# A cancer pharmacogenomic screen powering crowd-sourced advancement of drug combination prediction

**DOI:** 10.1101/200451

**Authors:** Michael P Menden, Dennis Wang, Yuanfang Guan, Mike J Mason, Bence Szalai, Krishna C Bulusu, Thomas Yu, Jaewoo Kang, Minji Jeon, Russ Wolfinger, Tin Nguyen, Mikhail Zaslavskiy, AstraZeneca-Sanger Drug Combination DREAM Consortium, Sock Jang, Zara Ghazoui, Mehmet Eren Ahsen, Robert Vogel, Elias Chaibub Neto, Thea Norman, Eric KY Tang, Mathew J Garnett, Giovanni Di Veroli, Stephen Fawell, Gustavo Stolovitzky, Justin Guinney, Jonathan R. Dry, Julio Saez-Rodriguez

## Abstract

The effectiveness of most cancer targeted therapies is short lived since tumors evolve and develop resistance. Combinations of drugs offer the potential to overcome resistance, however the number of possible combinations is vast necessitating data-driven approaches to find optimal treatments tailored to a patient’s tumor. AstraZeneca carried out 11,576 experiments on 910 drug combinations across 85 cancer cell lines, recapitulating *in vivo* response profiles. These data, the largest openly available screen, were hosted by DREAM alongside deep molecular characterization from the Sanger Institute for a Challenge to computationally predict synergistic drug pairs and associated biomarkers. 160 teams participated to provide the most comprehensive methodological development and subsequent benchmarking to date. Winning methods incorporated prior knowledge of putative drug target interactions. For >60% of drug combinations synergy was reproducibly predicted with an accuracy matching biological replicate experiments, however 20% of drug combinations were poorly predicted by all methods. Genomic rationale for synergy predictions were identified, including antagonism unique to combined PIK3CB/D inhibition with the ADAM17 inhibitor where synergy is seen with other PI3K pathway inhibitors. All data, methods and code are freely available as a resource to the community.

## Introduction

Personalized treatment with drugs targeted to a tumor’s genetics have resulted in remarkable responses, however patients often relapse. Multiple opportunities for drug resistance exist ^1^, beginning with the genetic, non-genetic and clonal heterogeneity inherent of advanced cancers, coupled with complex feedback and regulatory mechanisms, and dynamic interactions between tumor cells and their micro-environment. Any single therapy may be limited in its effectiveness, but drug combinations have the potential to overcome drug resistance and lead to more durable responses in patients. The molecular makeup of cancer cells and the mechanisms driving resistance will influence the optimal combination of mechanisms to target^1–3^.

High throughput preclinical approaches are crucial to determine and evaluate effective combination strategies. While empirical approaches are important for assessing the synergistic properties across drugs, the possible number of combinations grows exponentially with the number of drugs under consideration. This is further complicated by the complex disease and cellular contexts, rendering it impractical to cover all possibilities with undirected experimental screens ^4^. Computational approaches for predicting drug synergy are critical to guide experimental approaches for discovery of rational combination therapy ^5^.

A number of approaches have been developed to model drug combination synergy using chemical, biological, and molecular data from cancer cell lines ^6,7^ but with limited translatability to the clinic. A key bottleneck in the development of such models has been a lack of public data sets of sufficient size and variety to train computational approaches ^4,8,9^ particularly considering the diversity of biological mechanisms that may influence drug response. A further limit to the translatability of many published approaches is their reliance on data unavailable in a typical patient scenario, such as on-treatment tumor molecular profiles, and the use of opaque models which lack biomarker rationale for subsequent testing and diagnostic development.

To accelerate the understanding of drug combination synergy, DREAM Challenges partnered with AstraZeneca and the Sanger Institute to launch the AstraZeneca-Sanger Drug Combination Prediction DREAM Challenge. DREAM Challenges (dreamchallenges.org) are collaborative competitions that pose important biomedical questions to the scientific community, and evaluate participants’ predictions in a statistically rigorous and unbiased way, while also emphasizing model reproducibility and methodological transparency ^10^. This Challenge was designed to explore fundamental traits that underlie effective combination treatments and synergistic drug behavior. Specifically, it was structured to address the following translational questions using data available prior to drug treatment (mirroring a clinically relevant scenario to direct therapeutic choice): [i] how to predict whether a known (previously tested) drug combination will be effective for a specific patient, [ii] how to predict which new (untested) drug combinations are likely to yield synergistic behaviors in a patient population, and [iii] how to identify novel biomarkers that may help reveal underlying mechanisms related to drug synergy. We shared with the scientific community 11,576 experimentally tested drug combinations on 85 cancer cell lines, by far the largest open release of such data to date. Molecular data was provided for the untreated (baseline) cell lines, alongside chemical information for the respective drugs. Participants used the described data to train and test models, and were encouraged to extend computational techniques to leverage *a priori* knowledge of cellular signaling networks.

In this manuscript, we report on the results of this Challenge where we have identified novel and performant methods using a rigorous evaluation framework on unpublished data. Additionally, we describe the details of these approaches, as well as general trends arising from the meta-analysis of all submissions. Data, methods and scoring functions are freely available to benchmark future algorithms in the field. Finally we identify the mechanistic commonalities evident across predictive features used to reveal genomic determinants of synergistic responses, particularly between receptor tyrosine kinase and PI3K/AKT pathway inhibitors.

## Results

### 1. The largest public high-throughput drug combination screen covering diverse disease and target space

We collated a combinatorial drug sensitivity screen comprising 11,576 experiments each measured in a 6-by-6 dose matrix across 85 cancer cell lines (**Supplementary Fig. 1, Supplementary Table 1**). This dataset included cell viability response measurements to 118 chemically diverse compounds, and estimated synergy scores for 910 pairwise drug combinations with high reproducibility (**Supplementary Fig. 2**; see Methods). Information on the compounds included putative drug targets and, where available, their chemical properties. We also integrated deep molecular characterization of these same cell lines, including somatic mutations, copy-number alterations, DNA methylation, and gene expression profiles (**Fig. 1a-c**) measured before drug treatment ^11^.

**Figure 1:**
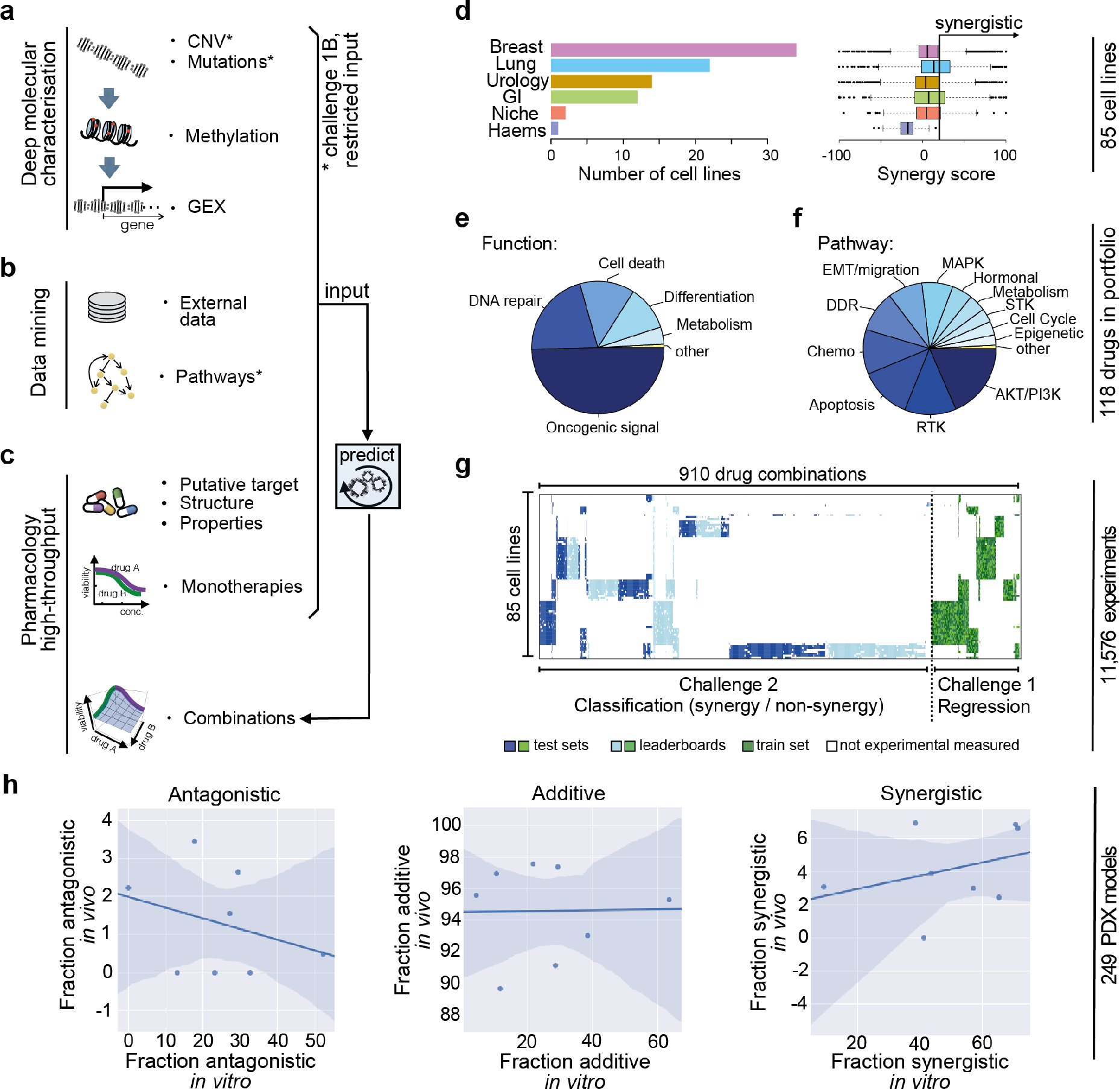
Drug combinations and cell lines profiled. (a) Molecular characterization of the cell lines included genetics, epigenetics and transcriptomics. (b) Participants were encouraged to mine external data and pathway resources. (c) Participants were provided the putative targets and chemical structures for ∼1/3 of cell lines to predict synergistic combinations. (d) The cell line panel contained 85 cell lines from 6 different cancer types. (e) The drug portfolio comprised approximately half oncogenic signaling targeting agents, and half cytotoxic compounds of which 14 were untargeted chemotherapies (f) Compounds split by the putative targeted pathway. (g) Sparse data was split into training set, leaderboard and independent test set for sub-challenge 1 and 2 and color coded accordingly, see legend in panel G. (h) Comparison of responses for overlapping combinations between in vitro (DREAM) and in vivo (Gao et al.) datasets shows translatability of synergistic combination-sample pairs (Pearson r = 0.34 for synergistic, r = -0.32 for antagonistic.)

The 85 cell lines are predominantly derived from tumors of the breast (N=34), lung (N=22), bladder (N=14), and the gastrointestinal tract (N=12) (**Fig. 1d**). Synergism for drug combination experiments were measured using the Loewe model, defined as increasing cell death beyond the expected additive effect of the individual compounds (see Methods). Drug synergy levels varied across disease types (**Fig. 1d**); in particular lung cell lines had over two-fold higher mean synergy than breast cell lines (p-value<7e-27). Of the 118 compounds tested, 59 were targeted therapies against components of oncogenic signaling pathways (see Methods), 15 of which target receptor tyrosine kinases (RTKs), 22 target PI3K/AKT signaling, and 9 target MAPK signaling (**Fig. 1e**). Across the pairwise drug combination experiments, 88% (N=797) of the unique pairs had drug targets within the same pathway and demonstrated markedly overall higher synergy levels (17.3 vs 7.3, p-value<2e-18) than the remaining 12% (N=113) whose drug targets were defined to be in different pathways. As part of the Challenge design, we ensured that drug targeted pathways and cancer types were proportionally distributed across sub-challenges and training/test data sets.

In order to assess the translatability of our *in vitro* combination screen, we collated response data for 62 treatments across ∼1,000 Patient-Derived Tumor Xenograft (PDX) models published by Gao *et al.* ^8^, which includes 24 drug combination pairs. Eight of these overlapped with our screen and were tested across 249 PDX models (**Supplementary Table 1**). We observed the highest correlation between *in vitro* and *in vivo* datasets (Pearson r = 0.34) for the fraction of samples with an observed synergistic response (see Methods), **Figure 1h**, suggesting confidence in the translatability of *in vitro* synergy response to *in vivo* PDX studies.

### 2. Comprehensive benchmarking of diverse computational prediction methods reveal accuracy of predictions reached the level of replicate experiments

The Challenge was divided into two primary sub-challenges. In sub-challenge 1 (SC1) participants were asked to predict synergy scores for drug combinations for which training data on those same combinations were available. In sub-challenge 2 (SC2), participants were asked to predict synergy status on drug combinations for which no training data was provided, thereby requiring participants to infer synergy using transferable data/knowledge patterns identified from previously seen independent compound pairs. SC1 was further subdivided into two parts: SC1A allowed the use of all available data for model prediction, while SC1B limited data use to just mutation and copy number variation (mimicking current clinical assay feasibility). A total of 969 participants of diverse geography and expertise registered for the Challenge (**Supplementary Fig. 3a,b**). 160 teams submitted across any portion of the Challenge and 78 teams submitted for final assessment. Specifically, SC1A received final submissions from 76 teams, 62 for SC1B and 39 for SC2. As scoring metric we used the average weighted Pearson correlation between predicted and known synergy values for SC1, and both the *-log_10_*(*p*) from a 3-way ANOVA and balanced accuracy (BAC) for SC2 (see Methods).

Across all teams, mean performance scores were R=0.24±0.01 and R=0.23±0.01 (weighted Pearson correlation ± standard error) for SC1A and SC1B respectively, and -log10(P)=12.6 (3-way ANOVA) for SC2. Despite the omitting several data types, teams performed only slightly worse for SC1B, Δprimary metric = 0.01 (P=0.90), compared to SC1A (**Fig. 2a**; **Supplementary Fig. 3c,d**). While teams employed many different methodological approaches to modeling drug synergy - including regression, decision trees, random forests, Gaussian processes, SVM, neural networks, text mining, mechanistic network-based and others (**Supplementary Fig. 4a**) - algorithm class showed little relationship to performance (**Supplementary Fig. 4b**). We observed that participants submitting to all sub-challenges rather than just one tended to do better (**Supplementary Fig. 3e**). The top winning team in all three sub-challenge was *Yuanfang Guan* (Y Guan) with primary metrics of 0.48, 0.45 and 74.89 in SC1A, SC1B, and SC2, respectively. Based on the primary metric in SC2, Y Guan performed considerably better (>5 Bayes Factor, based on bootstrapped metrics’ comparisons, see Methods) than other teams (**Fig. 2b**). All performance statistics and team rankings are available at the Challenge website (synapse.org/DrugCombinationChallenge).

**Figure 2:**
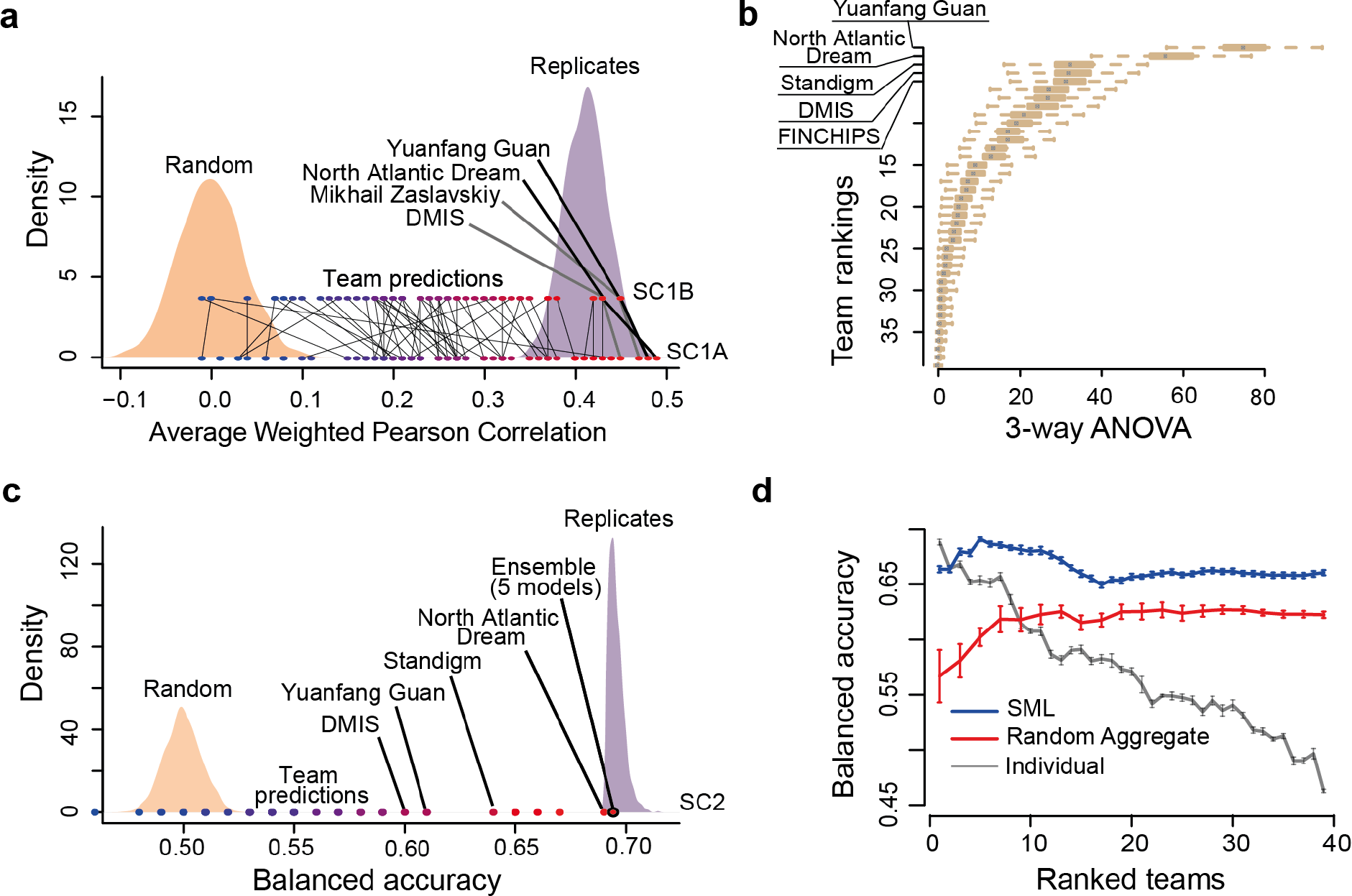
Performance of teams in the DREAM challenge. (a) Participant performance in SC1A and SC1B - the distribution of performance of random predictions was used to estimate a lower limit, and the distribution of synergy correlations between biological replicates were used to estimate the upper limit. (b) Participant performance ranked in sub-challenge 2 based on the primary metric, 3-way ANOVA. Distribution of bootstrap prediction performances for each team are shown by each boxplot with the dot showing their actual performance. (c) Participant performance plotted with upper and lower limits for SC2 based on the tie-break metric. Performance of random predictions were used to estimate the lower limit, and the performance of biological replicates were used to estimate the upper limit. (d) Ensemble models compared to the performance of individual models ranked from best to poorest performing in sub-challenge 2. SML is an ensemble of the best performing models based on estimation of their balanced accuracy. Random Aggregation is an ensemble combining a random combination of models. Standard error of mean represented by error bars are estimated from 10 random splits of the data.

To benchmark the performance of teams in the final rounds of SC1A/B and SC2, we established lower and upper bounds of performance. We defined the lower bound as the null model, *i.e.* random permutation of the synergy data across each cell line (see Methods). The upper bound was estimated as the level of correlation of synergy seen between experimental replicates. We observed that 83%, 85%, and 94% of submitted models performed better than random (5% FDR, see Methods) for SC1A, SC1B, and SC2, respectively. Team performances varied widely, but remarkably the top 15 models (20%) submitted to SC1A all reached a performance level comparable to the noise level observed in the experimental replicates (**Fig. 2a**), as did the top 13 models (21%) in SC1B. Proportionally fewer teams performed at the level of replicate experiments in SC2 based on the balanced accuracy (BAC), with North Atlantic Dream (NAD) coming closest to this bound (BAC=0.688; **Fig. 2c**).

Given the limited performance of SC2, we assessed whether an ensemble method - based on an aggregation of all submitted models - could yield a better overall model, a phenomenon called “wisdom of the crowd” ^10,12^. We used a Spectral Meta-Learner (SML) approach, and observed a marginal improvement in performance (BAC=0.693) over the best performing individual team (BAC=0.688) and an ensemble of any number of randomly chosen models (**Fig. 2d**). In SC2, SML ensembles including poorly performing models can achieve > 0.63 BAC.

### 3. Drug combination prediction is enhanced by leveraging biological relationships

Top performing teams (DMIS, NAD, and Y Guan) filtered cell line molecular features to leave only those in genes related to *a priori* cancer drivers (see Methods). These teams also consolidated pharmacological and/or functional pathway information associated with the molecular drug target, enabling one drug’s model to learn from data generated for another drug with the same target (Y Guan^16^ and NAD^14,15 16^).

To analyze each feature type’s importance, particularly whether incorporating molecular features and chemical/biological knowledge can increase prediction accuracy, we reengineered the DMIS and NAD models to use only cell line and drug labels as input features in SC1B (**Fig. 3**, see Methods). Using these models as baseline predictors, we were then able to iteratively substitute or add specific molecular features or external data sources (e.g. pathway/network information) to assess their importance in improving prediction (**Fig. 3a,b**). Surprisingly high primary metrics were found for the baseline model (**Fig. 3a**, 0.32) highlighting that drug and cell line labels alone hold predictive information. Drug target was the only feature to improve performance when swapped with drug or cell line labels (**Fig. 3a**, P=0.012), and removing both drug label and target resulted in the highest performance drop (**Fig. 3b**, -0.17). This result highlights the predictive value of the transferrable biological information encoded within drug target that is not available from unique drug labels. Mutational and copy number variation (CNV) data can similarly offer a barcode of cell identity to encode cell line label. However, where mutation data improved performance when replacing cell line label, replacement with CNV decreased performance significantly (**Fig. 3a**, P=8.8 × 10-6). Importantly, in all cases additional feature data increased performance when added to the baseline model, confirming that addition of biologically meaningful information truly adds to the model performance (**Fig. 3a**, P=0.009, 0.009, 0.002, 0.008, 0.021 adding drug target, 3 different pathways based and mutation features respectively). Ensembles of different feature sets improved prediction most when collectively increasing coverage of biological (pathway) complexity, leading to substantial increases in model performance (**Fig. 3a**, P=1.2 × 10-6).

**Figure 3:**
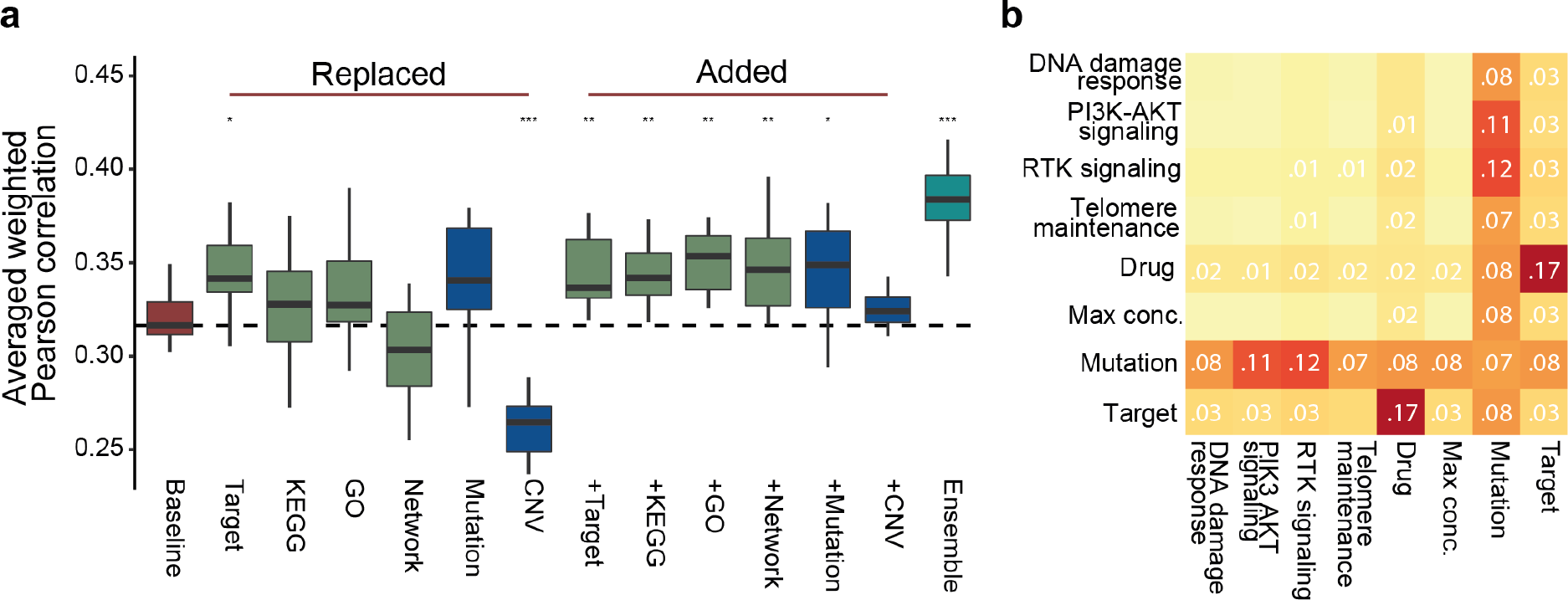
Feature impact. Drug target annotation is key in top performing algorithms, as is the meta information about variants including their functional impact and tumor driver gene status. (a) Cross validation based distributions of NAD primary metric of SC1B when replacing or adding drug/cell line label with respective features (baseline model has just drug and cell line label). *P<0.05, **P<0.01, and ***P<0.001 compared to baseline model (b) Heatmap of decrease in performance (average weighted Pearson correlation) of SC1B for DMIS support vector regression method when two feature types are removed at once (rows and columns).

### 4. A subset of combinations are poorly predicted by all teams, in-part explained by network connectivity of drug targets

While a global performance metric applied to all cell-lines and drug combinations provides a broad assessment of model prediction accuracy, we hypothesize that some models may be optimized for certain sub-classes of combinations and/or tumor types. We assessed the Pearson correlation between predicted and observed synergy scores for each combination in SC1A/B, and clustered teams by correlation of performance across combinations. Of the 118 combinations that had observed synergy scores >20 in more than one cell line, we identified 22 combinations predicted poorly by every participant (**Fig. 4a**, see Methods), and over 50 combinations were defined as well predicted across all teams.

**Figure 4:**
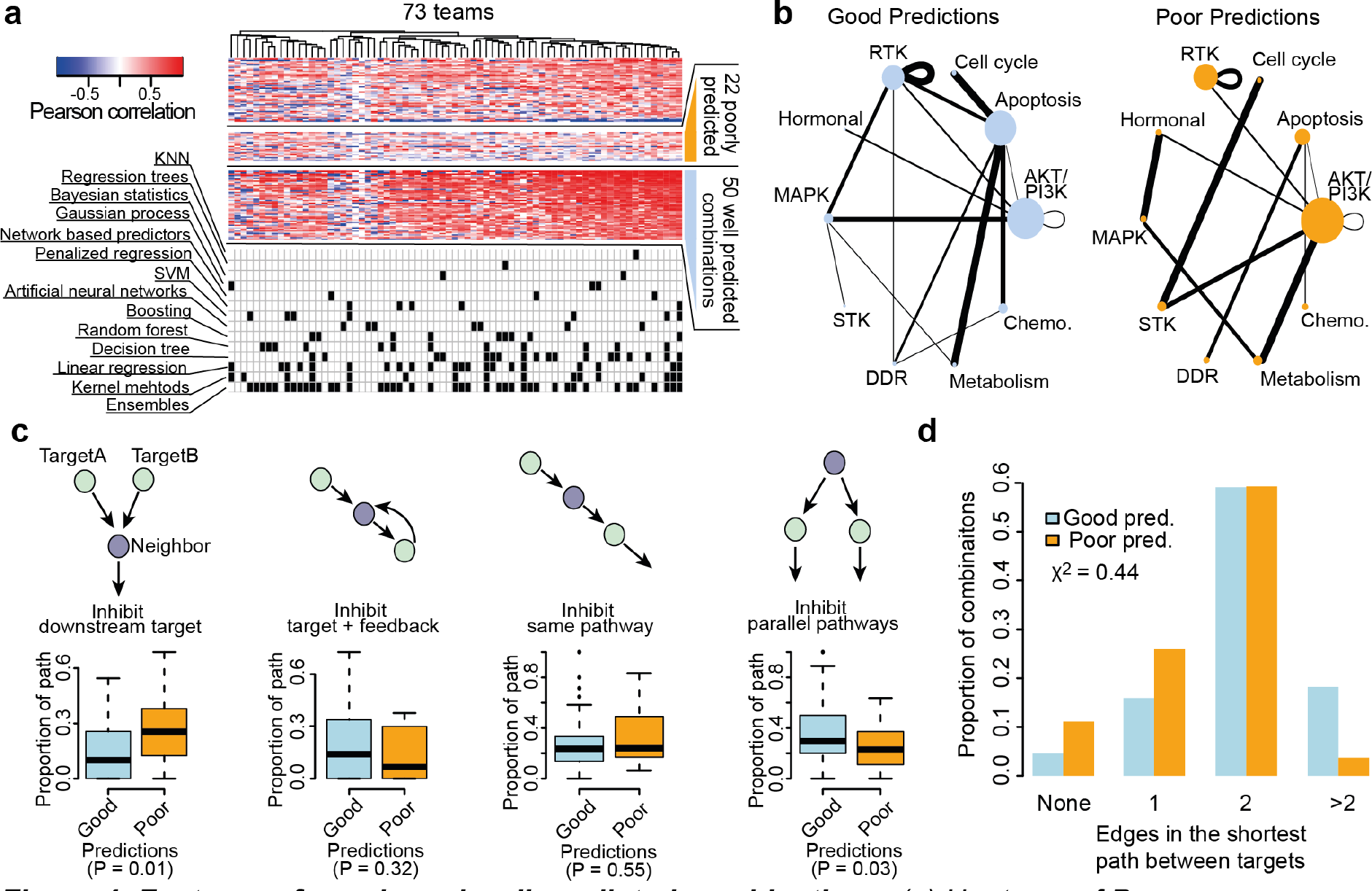
Features of poorly and well predicted combinations. (a) Heatmap of Pearson correlation between observed and predicted synergy scores for 118 combinations across 73 teams participating in SC1A/B. Algorithms used by each team is marked in the matrix below. (b) Combinations of pathways targeted. Size of node is proportional to number of drugs targeting specific pathway and width of edges is proportional to the number of drug combinations. (c) Types of interactions between the nearest neighbouring gene and the two drug targets of poorly and well predicted combinations. Boxplots show the difference in the proportion of interactions of each type for poorly and well predicted combinations (t-test). (d) Proportion of poorly and well predicted combinations for different network distances (minimum number of interactions in the OmniPath shortest path) between the two targets of a drug combination.

Surprisingly, neither the training data size per combination nor experimental quality showed notable difference between these universally poor and well predicted combinations (**Supplementary Fig. 5**). Higher performance (Fig. 4b, average Pearson correlation 0.37 vs 0.25; P=0.008) was observed for combinations inhibiting the PI3K/AKT pathway together with MAPK pathway or apoptosis pathway with either metabolism, cell cycle or receptor tyrosine kinases. Assessment of the interactions between drug targets and neighbouring proteins from *OmniPath*, a comprehensive compendium of literature-based pathway resources ^17^, revealed no differences in the somatic alteration frequency for targets or their first neighbors between the poorly and well predicted combinations (**Supplementary Fig. 6a,b**). We did observe a significant enrichment of well predicted combinations where both drugs’ respective targets were downstream of a common neighbouring protein (**Fig. 4c**, P=0.01), and conversely, we observed an enrichment of poorly predicted combinations where targets were both upstream (**Fig. 4c**, P=0.03). There was no significant difference (Chi-sqr P=0.44) in OmniPath protein network distance between targets of well and poorly predicted combinations (**Fig. 4d**).

### 5. Biomarkers of drug combination synergies

A typical shortfall of many machine learning algorithms is the lack of feature interpretability and experimentally testable logic-based rules. We took two approaches to identify biomarkers that may be predictive of drug synergies: a direct survey of participants through which predictive features were nominated for each drug pair (**Supplementary Table 2**); and retrospective work focusing on results from two of the best performing teams, NAD and DMIS, to deconvolute features most impactful to model predictions (**Supplementary Fig. 7, Supplementary Table 3**).

The survey-submitted biomarker results varied in detail and depth (**Supplementary Table 2**), but common genetic markers were apparent across good predictions in SC1B, including *EGFR, ERBB2, PIK3CA, PTEN, TP53* or *RB1*. Synergy or lack of synergy was commonly assigned to compound pairs targeting directly down-or up-stream of a mutated, amplified, overexpressed or deleted tumor drivers, including several well-established drug resistance markers. This enrichment suggests a hypothesis that monotherapy resistance biomarkers may increase the likelihood of synergy from combination with a second compound that can overcome that resistance. To systematically test this, we focused on a short list of tumour driver biomarkers (see Methods) and bootstrapped the significance with which they associated to resistance for each monotherapy. We applied a sliding threshold to this significance (see Methods) assessing the change in proportion of combinations with synergy if at least one compound succeeds the respective significance of association between a biomarker and monotherapy resistance (**Figure 5a**). We observed a significantly increased likelihood of drug synergy when subsetting for monotherapy resistance markers (Pearson correlation = -0.90, P=4.09 × 10^-38^). Furthermore, and emphasizing the *in vivo* translatability of these data, we observed the same trend in patient derived xenograft (PDX) models (**Fig. 5b**, Pearson correlation = -0.95, P=2.2 × 10^-49^). This observation supports the notion that drug resistance may be overcome by smart drug combinations targeting the putative resistance mechanism.

**Figure 5:**
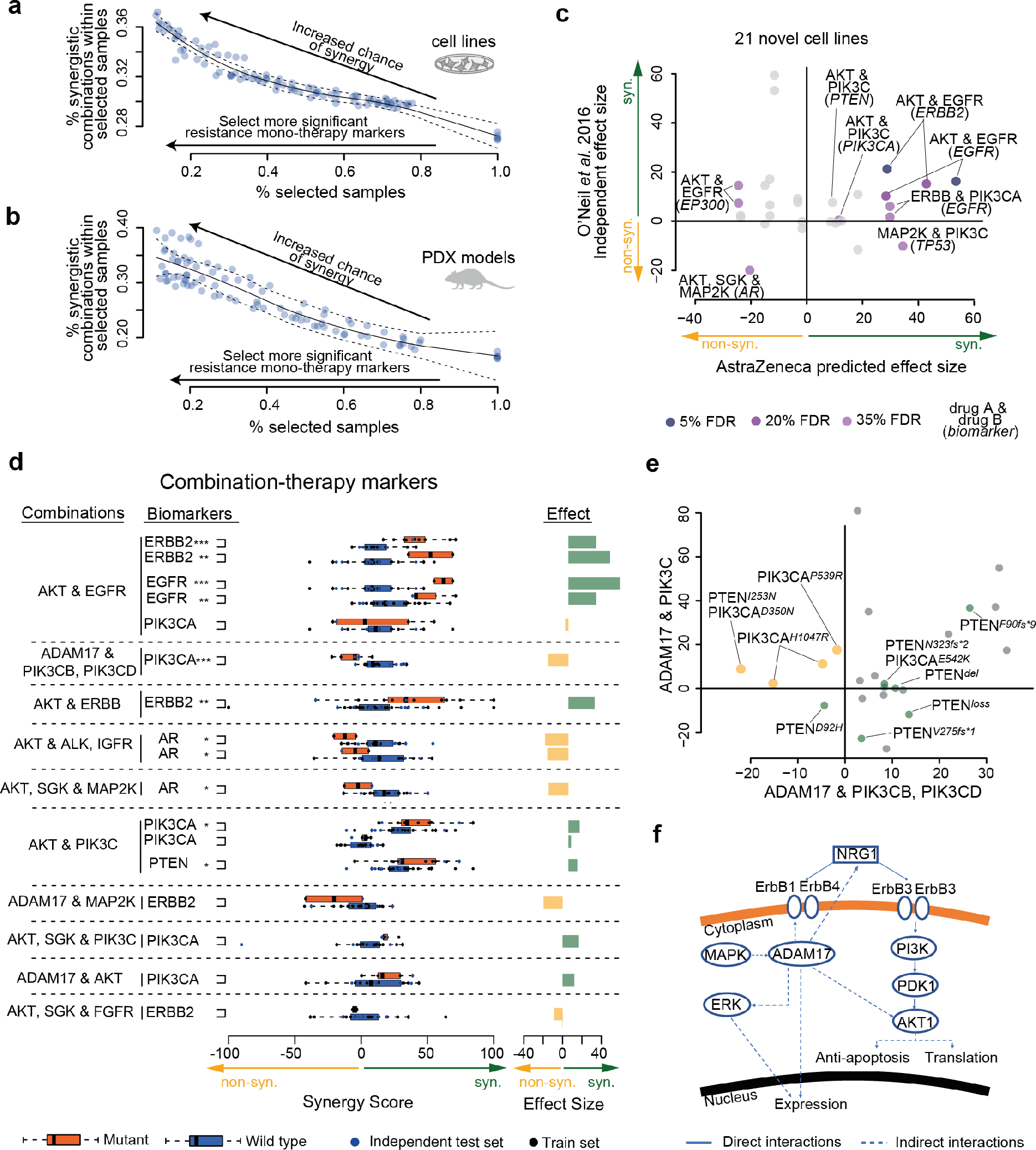
Post-hoc analysis of putative synergy biomarkers. (a) Cell lines and (b) PDX models [8] show increased frequency of synergistic drug combinations if they contain biomarkers with stronger association to monotherapy resistance. (c) Validation of biomarker predictions in new cell lines and independently screened drug combinations by O’Neil at al. 2016. (d) Synergy markers suggested by DMIS and NAD, when focusing on top weighted features from predictive models filtered for biological relatedness to drug targets, ‘***’=5%, ‘**’=20% and ‘*’=35% FDR. (e) Comparison of ADAM17 combined with PIK3CB/D against ADAM17 in combination with pan-PI3K3C inhibitor. (f) Network cartoon of PI3K signaling and role of ADAM17.

We also explored models of best performing teams and their chosen features, focusing on biomarker associations aligned to combinations for which the respective team had achieved a robust prediction accuracy (Pearson correlation > 0.5), with particular interest in the genetic biomarkers revealed through SC1B. Multiple validation criteria for quality, independency and reproducibility (see Methods) ^4,8,9^ were then applied to prioritize 13 feature-to-combination associations (**Fig. 5d**, **Supplementary Table 3**) for in-depth characterization of associated rationale, 7 associated with synergy and 6 with non-synergy.

Amongst the prioritized feature-to-combination associations were several genetic variants associated with synergistic responses to the combination of receptor tyrosine kinase (RTK) inhibitors with inhibitors of the downstream PI3K/AKT pathway. Amplifications or activating mutations in *EGFR* or *ERBB2* consistently predicted synergy from RTK + PI3K/AKT pathway inhibition across multiple independent drugs and data sets (**Fig. 5c,e**). Less direct relationships were also observed including combined AKT inhibition with EGFR inhibition in the *ERBB2* mutant setting or FGFR inhibition in the *EGFR* mutant setting. In addition to earlier observations, *EGFR* and *ERBB2* mutations were predictive of respective monotherapy responses (**Supplementary Fig. 8**), indicating that off-target effects are unlikely despite kinase domain homology. Combinations inhibiting multiple points within the PI3K/AKT pathway also showed synergy in the presence of upstream activation from mutations in *PIK3CA* or deleterious events in *PTEN* (**Fig. 5c**). Inhibition of the metalloproteinase ADAM17, known to influence RTK activity ^18^, also showed synergistic responses in a common subset of cell lines when combined with inhibitors of PI3K, AKT or MTOR, with a notable exception of antagonism unique to PIK3CB/D selective inhibition in *PIK3CA* mutant cell lines (**Fig. 5e,f**). Amplification and activating mutations in Androgen Receptor (AR) were also found to be associated with antagonistic effects for combinations targeting AKT and several MAPKs or RTKs, particularly MAP2K and IGF1R inhibitors (**Fig. 5c**).

### 6. Translatability - synergy and biomarker predictions from top performing teams generalize to independent data

We assessed the performance of top performing DREAM models on a smaller published screening experiment from O’Neill *et al* ^4^. O’Neill *et al* applied a different measure of cell death compared to the DREAM drug screens (Cell Titer-Glo vs Sytox Green). A similar correlation was observed among technical replicates in the O’Neill *et al* data set (rho=0.63) compared to the AZ-DREAM data (rho=0.56), however there was lower dispersion of synergy scores (**Supplementary Fig. 2c,d**) and fewer instances of extreme synergy scores in O’Neill *et al.*

Focusing on cell lines and drug combination tests (**Supplementary Table 1**) nonoverlapping between DREAM and O’Neill *et al* data, we observed that SC1A models from NAD and DMIS outperformed a random model for all new combinations in the O’Neill *et al* screen (**Fig. 6**, mean R = 0.07, P < 0.01). Interestingly, no substantial performance increase was observed when independent model predictions were made on mutational profile from the 10 cell lines in common between the two datasets, nor the 30 similar combinations with similar chemical properties. As in the main Challenge, combining these two models led to an improved prediction performance (**Fig. 6**).

In addition to the re-discovery of established and clinical drug combination biomarker relationships described above, we sought to systematically assess the reproducibility of biomarker predictions. NAD and DMIS explored a total of 509 genomic traits associated to drug combination synergies after respective pre-filtering (**Supplementary Table 3**) for SC1B. Features were ranked by their influence on model predictions for each of the well-predicted drug combinations (**Supplementary Fig. 7**, see Methods). We explored the top 5 ranked features for each well-predicted combination and consolidated drug-target centrically, giving a non-redundant list of 839 feature-to-combination associations. Filtering to results returned by multiple teams or with network/functional similarity between biomarker and drug target (see Methods) left 47 associations (**Supplementary Table 4**, 21 with FDR <35%). 7 of these associations could be mapped to independent cell lines in the O’Neil *et al*^4^ data set, with an overall Pearson correlation between DREAM and O’Neil effect sizes of 0.32 (Fig. 6b) illustrating the reproducibility of synergy markers.

**Figure 6:**
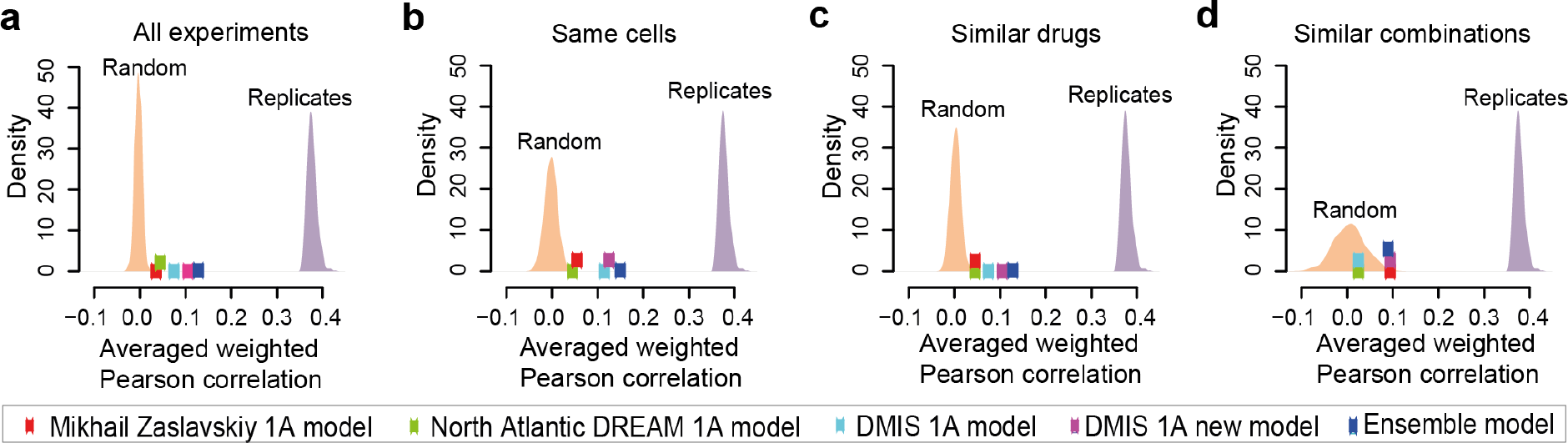
Translatability of top performing DREAM models to an independent screen by O’Neil et al ^4^. Performance of 1A models for predicting synergy scores in the O’Neil et al dataset by the best performing teams are plotted along with distributions of predictions from the random model and replicate experiments. Performance of predictions are shown for (a) all experiments in the O’Neil et al data set, and three subsets of the data set; (b) experiments that tested same cell lines as DREAM, (c) tested similar drugs as in DREAM (one drug in the combination with the same target), and (d) tested similar combinations as in DREAM (same targets for both drugs in the combination).

## Discussion

We have provided the largest unbiased *in vitro* drug combinatorial screen to date available to the scientific community. By demonstrating that trends represented in these data are reproduced *in vivo*, and that known *in vivo* clinically efficacious combination pairs can be identified, we offer confidence in the translatability of these data and their relevance for the prediction and characterisation of drug combination rationale. Our primary objective was to enable the development of a vast array of computational approaches to predict novel drug combinations, identify biomarkers for patient selection, and to comprehensively benchmark these approaches.

The results of the AstraZeneca-Sanger Drug Combination Prediction DREAM Challenge shed important light on the best strategies and limitations to predict drug synergy. By evaluating predictions from a large number of teams, we were able to uncover important strategies for predicting drug synergy from molecular and chemical traits. As with most DREAM Challenges, we observed that the machine learning method itself has little impact on overall performance, and that selective incorporation of biological knowledge can improve prediction accuracy. Aggressive pre-filtering that considers drug targets and gene relevance to cancer was successfully used by best performers to limit model complexity and to improve model generalizability. Despite the complexity of the problem, many teams achieved robust model performances, reaching the upper-bound of performance levels based on variability in experimental replicates. This was further confirmed when top performing models were applied to an independent data set, demonstrating robustness to assay variability, and context heterogeneity.

For pre-clinical data analyses, biological hypothesis discovery and mechanistic understanding is a more directly actionable goal than predictive accuracy. Models derived in pre-clinical data may prove the concept of predictability, hence our emphasis in SC1B to show prediction with data readily retrievable from a patient. However these models are unlikely to translate without further training in patient data since cell line panels do not comprehensively represent patient tumor characteristics. That said, predictive features and biological rationale revealed by these models can be directly tested and used to drive further research. We put special emphasis on incentivizing and retrieving this information, but found this challenging within a competition format that focuses on performance according to an objective scoring metric. In addition, accumulative small effects can explain good performance but are difficult to capture in post-hoc analysis with univariate test statistics. Given that this and prior DREAM Challenges indicate the machine learning method is less critical to performance than selection of biological features, we strongly advocate for the use of learners and mechanistic models with increased interpretability.

A comprehensive assessment of the predictive value of monotherapy was not completed in the Challenge format, in part due to initial miss-annotation of data, however retrospective analyses suggested it offered no significant improvement to well performing models (**Supplementary Fig. 9**). Despite minimal predictivity from monotherapy itself as a feature, resistance biomarkers predictive of monotherapy response do show predictivity of combination benefit. More synergy is also found where both drugs target downstream of a commonly interacting protein. Collectively these observations advocate for a more biologically rationalized approach, for example assembling a biomarker rationale by walking up- and downstream of the drug target to identify activated pathway components influencing monotherapy activity. Alternatively, more generic signatures of dynamic (e.g. transcriptional) output may first be used to identify a mechanistic rationale ^19,20,21,22^ to which causative genetic or epigenetic events can then be inferred and aligned as predictive features ^23,24^. A surprising result of our Challenge, however, suggested only modest improvement to prediction from inclusion of all data in SC1A compared to only genetics in SC1B.

A notable absence from the Challenge was the use of mathematical, boolean or logic based mechanistic pathway modelling approaches ^25–29^, likely due to the intensity of model creation. The dynamic nature of mechanistic models may offer an advantage by enabling consideration of the heterogeneity that exists across even apparently ‘clonal’ cell line populations ^21^. The increasing availability of published pre-derived mechanistic models for many cancer relevant pathways may soon make such an approach more viable. Given the strong benefit seen from inclusion of prior-knowledge, and as text based artificial intelligence technology matures, NLP and cognitive computational approaches to harness knowledge from world literature may also become of significant benefit.

Despite the limitations of the format, we were able to extract important insights to biomarkers for drug combinations. Given the dominance of RTK and PI3K/AKT pathway targeting agents in the Challenge data, it was not surprising that these revealed some of our strongest combination-feature relationships. In multiple cases this aligned to a two-hit hypothesis targeting the activating driver with a downstream pathway component. These included synergies between EGFR and AKT inhibitors in the presence of activating *EGFR* mutations ^30^, or AKT1/2 with pan-PI3K inhibitors in the presence of pathway activating or suppressing mutations in *PIK3CA* or *PTEN*, respectively. In some cases the biomarker rationale for AKT inhibitor synergy with RTK or MAPK inhibition was less direct and indicative of crosstalk and feedback signaling previously reported ^31^. Interestingly antagonism was observed in cell lines harboring activating mutations of *AR* ^32–35^. Feedback signaling resulting from AKT inhibition has been seen to drive AR activity which in turn can lead to the activation of the MAPK cascade ^35,36^, attenuating respectively targeting drug activity.

Synergy observed between ADAM17 and PI3K/AKT pathway inhibitors may work through independent inhibition of multiple cancer hallmarks, or via a more direct mechanism whereby inhibition of ADAM17 driven proteolysis and shedding of RTKs^18^ stabilizes and increases signaling through PI3K/AKT ^37,38^. Notably ADAM17 predominantly influences RTK’s other than EGFR/ERBB2^18^, and no benefit is seen in cells with mutations in these genes. Interestingly ADAM17 inhibition showed a unique antagonism with PIK3CB/D selective inhibitors within the *PIK3CA* mutant setting. Reduced synergy may result from a lessened dependency on PI3K paralogues in the presence of constitutively activated PIK3CA, or reduced benefit from ADAM17 loss in the extreme luminal/epithelial physiology of *PIK3CA* mutants. The apparent antagonism, however, suggests feedback following PIK3CB/D inhibition enhances mutant PIK3CA expression/activity. Indeed PIK3CB inhibition has been shown to result in elevated expression and activity of PIK3CA ^39^, and may also relieve the inhibitory effects of substrate competition or dimerization between PIK3CA and PIK3CB/D.

Looking forward, additional attention can be given to the one-fifth of of combinations poorly predicted by all teams in the Challenge. The rationale differentiating these combinations is non obvious, but data suggests some relationship to the complexity of network connectivity between drug targets and proximal biomarkers. Future Challenges should further address the question of how to optimize translation of preclinical results into the clinic ^40^. Where this Challenge addressed prediction of synergy for known drug combinations, an ability to predict truly novel beneficial drug combinations should also be explored. Most drug combinations effective in the clinic to date are effective due to the distinct effect of independent drugs on different subpopulations ^41^. Hence, identifying molecularly synergistic drugs, and how these affect inter- and intra-patient heterogeneity remains an essential area of future research. Furthermore the space of therapeutic combinations should be extended to include >2 drugs, covering targets in independent cell types such as subclonal tumor cell populations or cells of the tumor microenvironment and immune system^3^. These approaches can be complemented by adaptive and sequential strategies reactive to monitoring of the patient tumor and physiology. Success in these areas will be dependent on the availability and access to large-scale data needed for model development and validation. Public-private partnerships - as exemplified by this Challenge and the generosity of AstraZeneca to share their private data with the research community - will be critical to future efforts. We believe that the pharmaceutical and biotech industry will greatly benefit from these pre-competitive collaborations that accelerate basic research insights, and their translation into the clinic.

## Material and methods

### Drug combinations screening

All cell lines were authenticated at AstraZeneca cell banking using DNA fingerprinting short-tandem repeat assays and each bank is confirmed to be free from mycoplasma. Cells ordered from the global cell bank are cultured for up to 20 passages. Cell suspensions are counted using a haemocytometer and cells are re-suspended in full growth medium containing Pen/Strep to a final density for different cell line densities and for different seeding densities into 384 well cell culture plate. A volume of cells as determined by cell count and dependent on cell type was added to each well of a Greiner 384-well plate using a Multidrop Combi liquid handler and then incubated at 37C and 5% CO_2_ overnight in a rotating incubator. After seeding, plates were shaken to distribute the cells more evenly at the bottom of the wells and left to stand on the bench for 1hr to allow even settling of cells.

All plates were dosed with compounds solubilized in DMSO or PBS, or DMSO alone to provide comparable treatment and max control wells. Plates were dosed with compounds or DMSO only on an automated ECHO 555 acoustic reformatting system using the preconfigured DMSO and Aqueous calibration with DMSO normalized at final concentration of 0.14%v/v. After 5 days of incubation 5ul of 2uM Sytox Green working solution was added to each well of the 384-well plates (0.133uM final concentration) and the plates incubated for 1hr at room temperature. After incubation plates were read by the Acumen laser scanner to detect the number of Sytox Green stained cells. The total fluorescent intensity across the well was then read and the number of dead cells calculated by dividing this total fluorescence by the fluorescence of a single cell. The plates were re-read on the Acumen to give a total cell count. A live cell count was then determined by subtracting the dead cell count from the total cell count.

### Quantifying combination synergy and antagonism

Monotherapy dose-responses of each drug in a combination was modeled as a sigmoidal curve and fitted to a classical Hill equation. In order to identify synergy or antagonism, an additive effect was first derived based on single agent dose-response curves using the Loewe model (Fitzgerald 2006; Geary 2013). The Loewe model relies on the isobole equation which was solved numerically for all drug concentration values in order to calculate *A*(*a*,*b*) and then derive *S*(*a*,*b*)=*E*(*a*,*b*)-*A*(*a*,*b*). the synergy distribution *S*(*a*,*b*) was summarized d by integrating *S*(*a*,*b*) in logarithmic concentration space, what we called total synergy using Combenefit v1.31 ^42^

### In vitro - in vivo Translatability

Response data for 62 treatments across ∼1,000 PDX models were derived from Gao et al. [^8^]. Due to the anonymity of compound labels, primary targets from both datasets were utilised to identify overlap.

#### In vivo response class definitions

As synergy scores were not available for the Gao et al. dataset, ‘Best Response’ (Complete Response – CR, Partial Response – PR, Stable Disease – SD, Progressive Disease - PD) for each combination-PDX pair were extracted and compared with monotherapy ‘Best Response’ of each compound in the combination on the same PDX model. This was represented numerically where CR=4, PR=3, SD=2 and PD=1. Synergy was assigned to a change of +2 or more, and Antagonism to a change of -2 or less. A change of +1, 0 or -1 was assigned Additive, considering an element of experimental flexibility. Cases where best response has been observed as a range over time (PR->->PD), the earliest response was considered as we hypothesise this to reflect *in vitro* response in a more realistic sense for comparison.

#### In vitro response class definitions

Response scores defined by the Loewe synergy model were considered in ordered to define *in vitro* response classes. Synergism was defined as Loewe scores >= 5, Antagonism <= -5, and rest are classed as Additive.

### Molecular characterisation

The 85 cell lines were molecularly characterized, including:xs

1. Mutations from whole exome sequencing with Illumina HiSeq 2000 Agilent SureSelect (EGAS00001000978)
2. Copy number variants from Affymetrix SNP6.0 microarrays (EGAS00001000978)
3. Gene expression from Affymetrix Human Genome U219 array plates (E-MTAB-3610)
4. DNA methylation from Infinium HumanMethylation450 v1.2 BeadChip (GSE68379)

#### Mutations

Mutations were called with CAVEMAN ^43^ and PINDEL ^44^ as reported in ^11^. Variants were provided without further filtering, including putative passenger mutations, germline variants and potential cell line artefacts, which are in total 75,281 mutations in 85 cell lines.

#### Copy number events

Copy number variants (CNVs) are called with the PICNIC ^45^ algorithm using the human genome build 38 as the reference. CNVs might be wild type, deletion or amplification of certain segments in a chromosome. One or multiple genes can fall within such segments. We reported copy number for the major and minor allele on gene and segment level.

#### Gene expression

Gene expression was processed as described in ^11^ including Robust Multi-array Average (RMA) normalization with the R-package ‘affy’ (Gautier, Cope, Bolstad, & Irizarry, 2004). Gene expression for 83 cell lines across 17,419 genes (HGNC labels) was reported; no expression was available for MDA-MB-175-VII and NCI-H1437.

#### DNA methylation

We reported for 82 cell lines the *beta* and *M* values ^46^ for 287,450 probes; no methylation was available for the cell lines SW620, KMS-11 and MDA-MB-175- VII. In an additional processing step, CpG sites were compressed to CpG ilse with the definition from UCSC genome browser ^47^, resulting in a total of 26,313 CpG ilse based on either *M* or *beta* values.

### Drug properties

The identity of all compounds was anonymized, but for all agents the putative targets are revealed. The gene names of the protein targets are listed with ‘*’ denoting any character if the target is a protein family. Furthermore, for 58 compounds the Molecular weight, H-bond acceptors, H-bond donors, calculated octanol-water partition coefficient, Lipinski’s rule of 5, and their SMILES (Simplified Molecular Input Line Entry Specification) are provided. Drugs were grouped into pathways and biological processes manually according to their protein targets (**Supplementary Table 1**).

### Challenge organization

The Challenge consisted of 2 sub-challenges, each with multiple rounds: a leaderboard, validation, bonus and collaborative round. sub-challenge 1 had 4 leaderboard rounds that lasted 8, 6, 5, and 5 weeks, while sub-challenge 2 had 3 leaderboard rounds that lasted 12, 7, and 5 weeks. Participants were given a leaderboard dataset to build a model and generate 3 prediction files per leaderboard round. Scores were returned to participants so that they can improve their model throughout these rounds for their one submission to the final round which was scored against a held-out dataset. The final round lasted for 2 weeks which was then followed by a 9 week bonus round and 10 week collaborative round.

### Challenge pharmacology data splits

In sub-challenge 1, participants were asked to predict drug synergy of 167 combinations in the panel of 85 cell lines. The synergy data of each drug combination was partitioned into 3 sets: a training data set (3/6-50%), a leaderboard set (1/6-16.7%), and validation set (2/6-33%) of treated cell lines. sub-challenge 2 leveraged data for remaining 740 drug combinations not overlapping with those used in sub-challenge 1, although data for some of the same compounds (in combination with different compounds), homologous compounds (i.e. same target, but different chemical structure), and cell lines were included. A leaderboard set (370 combinations) and a final validation set (370 combinations) were randomly split, which are mutually exclusive from each other as well as from sub-challenge 1.

### Challenge Scoring Metrics

#### Sub-challenge SC1, Primary Metric

The primary metric was an average weighted Pearson correlation (*ρ_w_*) of the predicted versus observed synergy scores across each individual drug combination, *i*. The weight for a given drug combination *i* was 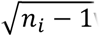 where *n_i_* is the number of cell lines treated with the drug combination. This resulted in the following primary metric for SC1A&B,

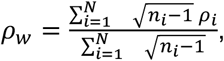

where *N* = 167were the tested drug combinations.

#### Sub-challenge SC1, Tie-Breaking Metric

The tie-breaking metric was identical to the primary metric except that it was applied to the subset of drug combinations that have at least one cell line with synergy score *S_ci_* ≥ 20 in the held-out test set (*S_ci_* = synergy score at cell line *c* and drug combination *i*). Neither the subset of drug combinations nor its size (*N* = 118) was revealed to participants prior to final evaluation.

#### Sub-challenge SC2, Primary Metric

The primary metric was a sequential three-way ANOVA, which tested the separation of held-out synergy scores by predicted synergy (= 1) and predicted non-synergy (= 0). In the sequential three-way ANOVA (type 1), we controlled for systematic drug and cell line effects, and evaluated variance explained by a given team’s synergy predictions. We define the primary metric as

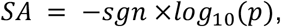

where *sgn* is the sign of the effect size (positive or negative separation by prediction), and *p* is the p-value (F-statistic) computed from the ANOVA distinguishing predicted synergy (= 1) from predicted non-synergy (= 0) across all experimentally measured synergy scores.

#### Sub-challenge SC2, Tie-Breaking Metric

As the tie-breaking metric, we used balanced accuracy (BAC) using discretized synergy scores *S_ci_* ≥ 20

#### Applying the Tie-Breaking Metric

In each sub-challenge, we estimated a Bayes Factor (BF) using a paired bootstrapped approach to determine whether a team’s score was statistically indistinguishable from another. In the event that a team’s scores were determined to be statistically equivalent, we then applied the tie-breaking metric. To estimate the BF, we sample with replacement from the *M* observations of the given sub-challenge and computing the primary metric (pm) for each team 1000 times. For a given team, T, *K_T_* was computed by

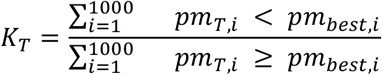

Where *pm_besti_* is the bootstrapped primary metric at iteration *i* for the team with the highest primary metric (non-bootstrapped).

### Assessing performance of individual combinations

Combinations defined as poorly predicted had an average predicted vs observed Pearson correlation across teams in the range seen with a random predictor (**Supplementary Fig. 8**, -0.25 and 0.25). In contrast, well predicted combinations had an average Pearson correlation across teams of above 0.5.

### Independent validation on O’Neil *et al* Merck screen

In order to assess the utility of features and the predictability of the learning algorithms in new contexts, we provided the participants an independent large-scale oncology combination screen published recently ^4^. The O’Neil *et al* dataset consists of 22,737 experimental endpoints covering 583 doublet combinations across 39 diverse cancer cell lines. 38 experimental compounds and approved drugs were included in this combination screen using a 4-by-4 dosing regimen. Raw cell viability measures for each combination experiment were processed through Combenefit ^42^ and dose response surfaces were tested against the Loewe synergy model (same as in the Challenge). While there are 6 approved drugs, 49 targets, and 10 cell lines in common between the Challenge and O’Neil *et al* datasets, the total number of exact experiments (Compound A – Compound B – Cell line) overlapping is below 100, giving the participants a highly independent validation set for their prediction algorithms. This information was provided to best performing teams in the Challenge, along with a dictionary of curated chemical structures and putative targets for each. Prediction models were trained on the released Challenge dataset and made synergy score predictions on the O’Neil *et al* dataset. Metrics for SC1 and SC2 were used to assess prediction performance.

### Individual Prediction Models

Full description and implementation of models used by teams in the final submission to DREAM can be downloaded from:
www.Synapse.org/AstraZeneca_Sanger_Drug_Combination_Challenge_Leaderboards. Top performing prediction models in SC1 and SC2 made use of genetic features relating to the gene targets of the drugs. Feature selection from the models enabled nomination of putative biomarkers for drug combination synergy (see Supplementary Material).

### Ensemble Models

Sub-challenge 2 participant models were aggregated using two types of ensemble models Spectral Meta-Learner (SML) and Random Aggregation. SML choses predictions from *n* methods to aggregate based on an estimation of balanced accuracy for each method without using actual labels ^13,48^. Random Aggregation is the traditional way that people aggregate models by giving equal weight to each method. We randomly pick *n* methods (do this 10 times) and for *n* methods we compute the average balanced accuracy and the error.

### Monotherapy Biomarkers and Synergy enrichment

Monotherapy markers are the mutational status of genes, either mutated or copy number altered, from the pan-cancer binary event matrix (BEM) ^11^, which separate the monotherapy response into sensitive versus non-response. The likelihood of separation was estimated with a Wilcox Rank Sum test. From most significant monotherapy marker to lowest in 0.1 steps of -log10(p-value), we accumulative evaluated the percentage of synergistic combinations with at least one monotherapy marker. This analysis was bootstrapped 10 times with 80% of the pharmacology data.

### Synergy Biomarkers

A short list of putative synergy biomarkers were derived from the 5 highest ranked features of well predicted drug combinations (Pearson > 0.5) from the two best performers NAD and DMIS. Features were ranked based on their feature weight or importance for given well predicting model. This gene-to-combination short list, was filtered for associations predicted by both teams, or genes biological related to the drug target defined as either the gene being the target itself, a short distance to it in OmniPath signaling network (2 molecules up- or downstream) or GO term similarity ^49^ larger than 0.7. This resulted in a list of 47 gene-to-combination associations that we further studied. A gene within this list is considered mutant if it was deleted, amplified (more than 7 copies) or mutated in any sense, resulting in an extended BEM ^11^. We calculated the p-value for each suggested association with an ANOVA correcting for tissue of origin and multiple hypothesis testing via Benjamini Hochberg. The effect sizes is the mean difference in synergy score between mutant and wild type cell lines.

For external validation of those putative biomarkers of synergy, we focused on drug combinations in O’Neil *et al.* 2016 ^4^, ALMANAC ^9^ and additional experimental data from AstraZeneca (**Supplementary Table 3**). We validated biomarkers in two different contexts, (i) for cell lines overlapping with DREAM, considered as biological replicates, and (ii) cells non-overlapping for predictions on novel cell lines.

Literature evidence for the shortlisted combination-biomarker associations was identified through PubMed search. The aim was to identify published evidence of (i) the combination-biomarker association, (ii) the combination but not the specific biomarker, and (iii) either one of the targets and the biomarker association. The publications were further categorized into *in vitro, in vivo*, and preclinical studies. Publications that discuss the specific combination-biomarker association have been highlighted in red (**Supplementary Table 4**). In summary, synergy biomarker were derived from best performer models, and highlighted based external validation as well as literature support.

### Accession codes

Full description of generation methods provided to all participants in this Challenge can be downloaded from https://www.synapse.org/DrugCombinationChallenge, while full data is available from https://openinnovation.astrazeneca.com/data-library.html.

## Acknowledgements

We thank the Genomics of Drug Sensitivity in Cancer and COSMIC teams at the Wellcome Trust Sanger Institute for help with the preparation of the molecular data, Denes Turei for help with Omnipath. Funding from the European Union Horizon 2020 research and innovation program under grant agreement No 668858 PrECISE to JSR.

## Authors contribution

MPM, DW, TY, ISJ, TNo, GYD, SF, GS, JG, JRD and JSR designed the challenge. The top-performing approach was designed by YG. Data analysis for the top-performing approach was conducted by MPM, DW, MM, BS, KCB, JK, MJ, RW, TNg and MZ. The DREAM Consortium provided drug synergy and biomarker predictions, as well as method implementations and descriptions. MPM, DW, MM and TY performed analysis of challenge predictions. MPM, DW, MM, BS and KCB interpreted the results of the challenge and performed follow-up analyses for the manuscript. EKYT, MJG and SF generated experimental data. MPM, DW, YG, MM, BS, KCB, TY, JK, MJ, RW, TNg, MZ, DREAM Consortium, ISJ, TNo, EKYT, MJG, GYD, SF, GS, JG, JRD and JSR wrote the manuscript. JG, JRD and JSR supervised the project.

